# Chemogenetic Manipulation of the Subthalamic Nucleus-Substantia Nigra Pars Reticulata Pathway Promotes Recovery in HemiParkinsonian Rat Models

**DOI:** 10.1101/2025.08.28.672861

**Authors:** Nassim Stegamat, Rupert Smit, Jacquelynn Rajavong, Thomas Campion, Sraavya Pinjala, George Smith

## Abstract

Motor control through the basal ganglion is mediated primarily thought two pathways, the direct and indirect pathways. The indirect pathway functions in opposition to the direct pathway, balancing excitatory and inhibitory control over motor movement. In Parkinson’s disease, the indirect pathway is thought to suppress movements through excessive excitation of inhibitory neurons within the globus pallidus internus and substantia nigra pars reticulata. In an attempt to restore this balance, we induced the expression of GAD65 in glutamatergic subthalamic neurons to enable the co-release of GABA. Previous studies indicated co-expression of GABA with glutamate reduces glutamate excitability and might restore balance to the network. Indeed, we observed a reduction in post-lesional amphetamine-induce rotations in 6OHDA lesioned rats co-transduced with GAD65 and excitatory DREADDs and activated by CNO. This study indicates rebalancing inhibitory and excitatory output from the subthalamic nucleus can have a positive effect on motor outcome.

## Introduction

Parkinson’s Disease (PD) is the second most common neurodegenerative disease in the elderly population, with a higher prevalence in men, independent of race and social class; it affects approximately 1.5 to 2.0% of the elderly population over 60 years of age and 4% of those over 80 years of age.(1). PD is caused by the necrosis of dopaminergic neurons in the substantia nigra pars compacta (SNc). This ultimately results in an overall decrease in the neurotransmitter dopamine in the synaptic cleft. The monoamine oxidase B (MAO-B) degrades dopamine, promoting glutamate accumulation and oxidative stress with the release of free radicals, causing excito-toxicity(1). PD symptoms manifest as progressive physical limitations such as rigidity, bradykinesia, tremor, postural instability and disability in functional performance(2). Considering that there are no laboratory tests or biomarkers to confirm the disease, the diagnosis of PD is often made clinically by analyzing the motor features. There is no cure for PD, and the pharmacological treatment consists of a dopaminergic supplement with levodopa, COMT inhibitors, anticholinergics agents, dopaminergic agonists, and inhibitors of MAO-B, which basically aim to control the symptoms, enabling better functional mobility and increasing life expectancy of the treated PD patients(1).

For Parkinson’s patients who are unable to maintain appropriate quality of life with pharmacologic agents, deep brain stimulation (DBS) is an option where an electrode is implanted into the subthalamic nucleus (STN) of the brain then powered by a battery pack implanted sub-clavicularly(3).

This treatment modality is based on PD’s augmentation of the excitatory and inhibitory circuits in basal ganglia pathways. Under normal brain physiology, basal ganglia regulates movement through two pathways that work in opposition, the direct and the indirect pathway(4). In PD, the STN becomes over-excited augmenting the indirect pathway causing excessive suppression of motor activity due to the imbalance of direct and indirect pathway input on the primary motor cortex. Input to the motor cortices is transmitted from the STN initially through its projections to the globus pallidus internus (GPi) and substantia nigra pars reticulata (SNr)(5-7). DBS is able to recover proper motor function by suppressing the STN’s over-excitation and restoring equilibrium amongst the basal ganglia motor pathways. While DBS is a widely accepted treatment for PD, the invasive nature of the procedure entails various complications including intracranial hemorrhage, implant infection, electrode corrosion, cognitive dysfunction, etc. Furthermore, battery packs implanted sub-clavicular need to be replaced every 3-5 years requiring further surgery down the road(8).

Advancements in PD research have progressed to investigating less invasive treatments to model DBS including gene therapy which has been a promising avenue. In a 2002 study, Dr. Luo and colleagues modeled DBS chemo-genetically by injecting adeno-associated virus encoding glutamic acid decarboxylase (GAD65) into the STN of a Parkinson’s induced rat(9). The transduction of GAD65 into glutamatergic neurons allowed for the release of both GABA and glutamate to reduce excitation of post-synaptic neurons. This reduced the indirect pathway excitation into the GPi and SNr, ultimately remediating parkinsonian behavior. Further advancements in genetic therapy included the invention of designer receptors exclusively activated by designer drugs (DREADDs) which act as synthetic neurotransmitter receptors to drive inhibition or activation of the basal ganglia at will via stimulation by a synthetic ligand (clozapine-n-oxide, CNO)(10). This technology was explored in macaque monkeys where their STN was selectively inhibited using DREADDs. They found increased neural spike variability when the STN was inhibited demonstrating the efficacy of DREADDs and further substantiating the STNs role in motor output organization(11).

Our project aims to further advancements in PD research by modelling DBS in a less invasive fashion through a combination of GAD and DREADDs genetic therapeutics. With this combination, we intend to convert excitatory neurons in the STN into inhibitory neurons via GAD expression, then excite the now inhibitory STN via excitatory DREADDs to reduce the indirect pathway’s excessive suppression of motor activity in PD and recover function.

## Results

To phenotypically alter the neurotransmitter specificity of subthalamic neurons, we used intersectional genetics. This was accomplished by injecting retrograde transportable AAV encoding either GAD65, hM3Dq or both into the SNr and AAV encoding CreR into the STN (Figure 1). For histological analyses the GAD65 virus co-expressed GFP (green), whereas, the hM3Dq virus co-expressed mCherry (red). Analyses of the STN showed good labeling of neurons for both red (Fig. 2A) and green (Fig. 2B) fluorescent markers, with about 80% showing both marker proteins (Fig 2C). Within the SNr we observed mostly subthalamic GAD 65 axonal labeling (Fig. 2E) with some red neuron labeling (Fig. 2D). In the SNr we were able to also visualize a minimal amount of co-expression of hM3Dq neurons with GAD65 (Fig. 2F).

**Figure 1.**
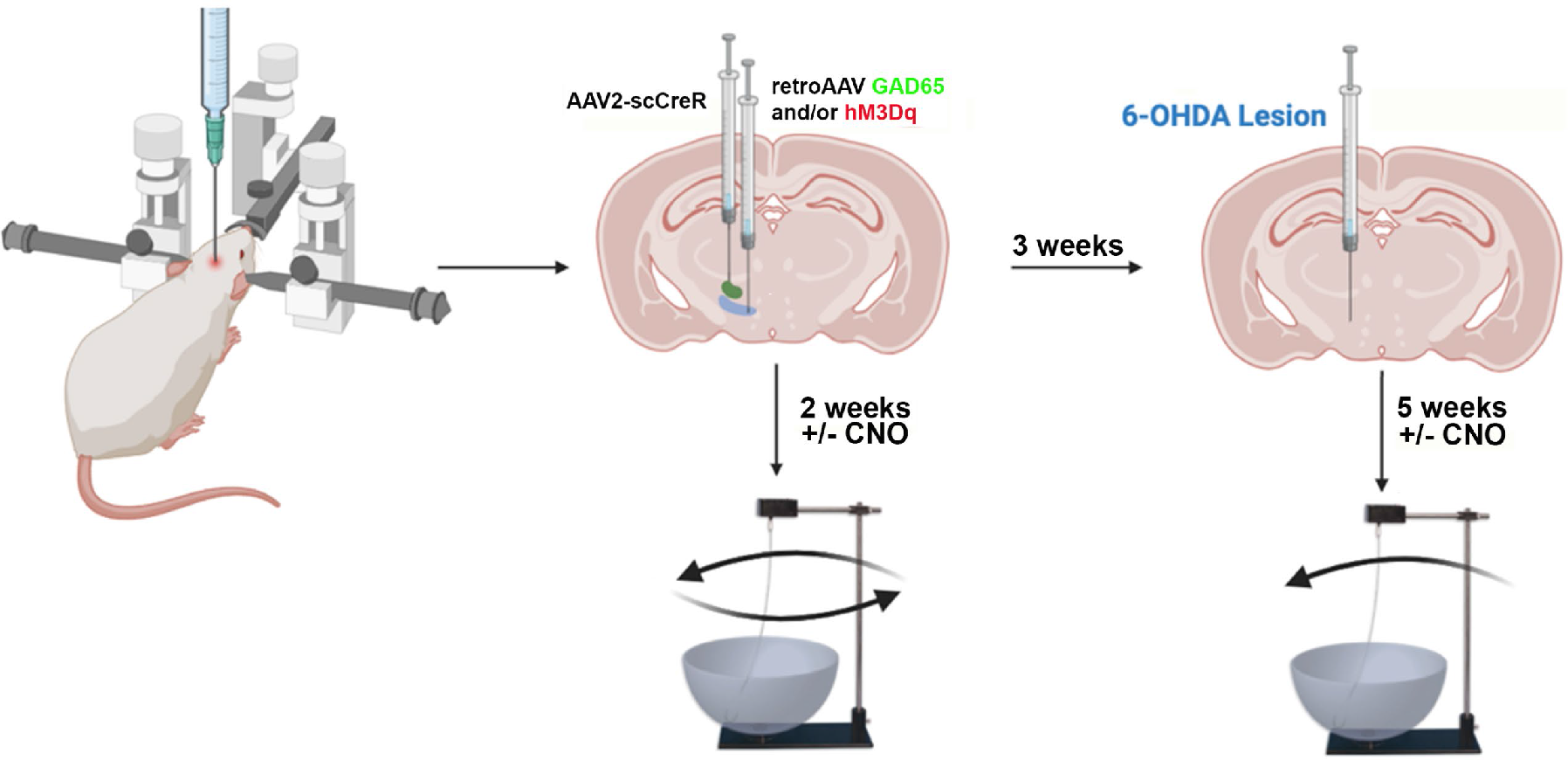
Illustration showing stereotaxic injections of viruses and 6OHDA into the STN/SNr and medial forebrain bundle respectively. Rats were placed in a stereotaxic device for injections. The SNr was injected with either AAV-retro-hSyn-DIO-GAD65-P2A-eGFP, AAV-retro-hSyn-DIO-hM3Dq-mCherry or both (1ul total) and the STN injected with AAV2-scCre (0.2 ul total). One weeks later rats were examined for amphetamine-induced rotation with and without CNO. The following week 6OHDA was injected into the medial forebrain bundle to kill dopaminergic neurons within the SNc. After 5 weeks post lesion, rats were retested for amphetamine-induced rotation with and without CNO.

**Figure 2.**
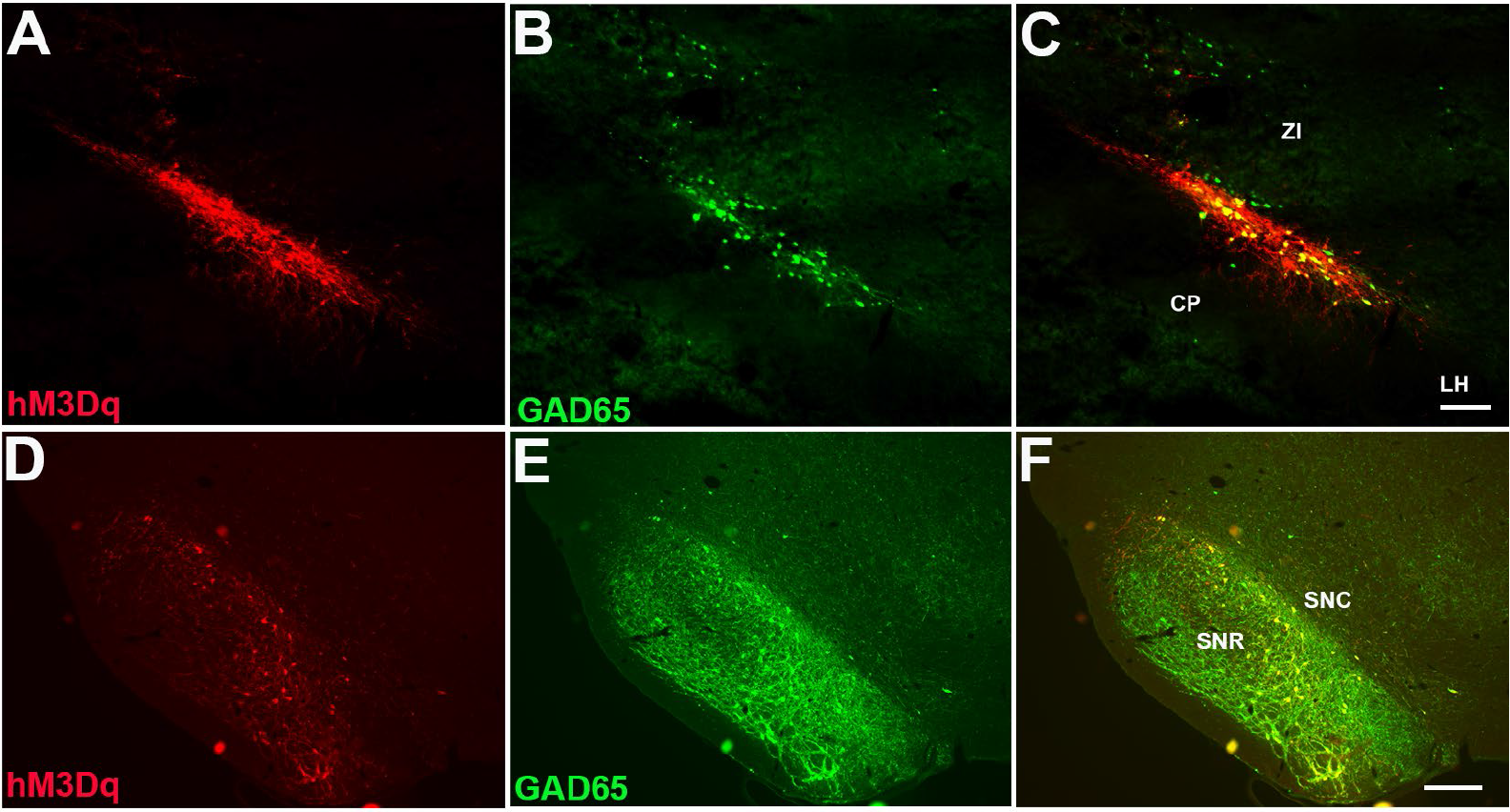
Expression of transgene fluorescent marker proteins in either the STN (A,B,C) or the SNr (D,E,F). Within the STN most neurons showed co-labeling for hM3Dq (red, A,C) and GAD65 (green, B,C). Within the SNr there were numerous GAD65 expressing axons with a few interspersed GAD65 + hM3Dq co-labelled neurons(F). Scale bars 200 um (A,B,C) and 250 um (D,E,F). ZI – zona inserta, CP – cerebellar peduncle, LH – lateral hypothalamus, SNR – substantia nigra pars reticulata, SNC - substantia nigra pars compacta.

To determine if altering these glutamatergic neurons to also release GABA acted to reduce movement disorders in rats with unilateral Parkinson’s like disorder, we injected 6OHDA into the left medial forebrain bundle. This procedure effectively destroys dopaminergic neurons within the left SNc causing the animal to turn ipsilateral to the lesion (counterclockwise, CCW) in the presence of amphetamine induction. All 6OHDA post-lesional rats irrespective of treatment showed a significant increase in the number of CCW rotations after amphetamine administration (Fig. 3A). Rats demonstrating a recovery in Parkinsonian phenotype would spin in a clockwise (CW) fashion or the rats would have an overall reduction in CCW rotations. This is due to the fact that rats with a lesioned basal ganglion will have deficiencies contralateral to that basal ganglia; therefore, the rats right side would be affected in our experiment forcing the rat to use their left side and only allowing for CCW rotations. The therapy acts to remediate the effects of the lesion and allow the rat to use both paws more evenly; thus, rats would be trending towards an increase in CW rotations and a decrease in CCW rotations.

**Figure 3.**
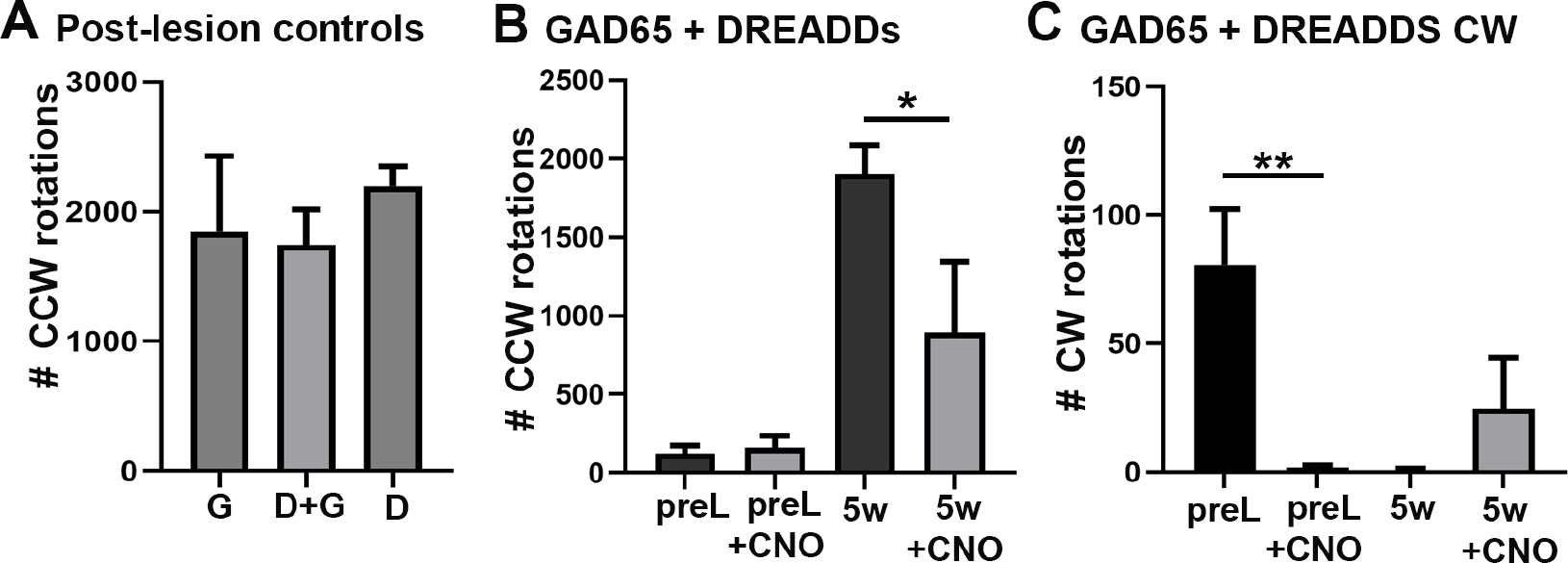
Changes in amphetamine-induced rotations with co-expression of both GAD65 and hM3Dq. A) Five weeks after 6OHDA lesion, rats in all groups showed similar levels of counterclockwise rotations. (B) Treatment with CNO reduced the number of counterclockwise rotations in GAD65/hM3Dq expressing rats. (C) Treatment with CNO increased the number of clockwise rotations in GAD65/hM3Dq expressing rats. * < 0.05, **< 0.01

Here the therapies would be activated upon CNO stimulation, each rat served as their own control and were compared to themselves with or without CNO stimulation at the same time point. Prior to 6OHDA lesioning we tested the rats receiving AAV for both GAD65 and hM3Dq with CNO and observed a slight but statistically significant decrease in the clockwise rotations using a two-tailed paired sample t-test to compare values between pre-lesioned rats with and without CNO (Fig 3 C). Pre-lesioned rats without CNO stimulation had a higher mean CW rotation (M = 80.7 ± 36.9) compared to rats with CNO stimulation (M = 1.7 ± 1.5), with a mean paired difference of 79.0 ± 39.0. Post-lesioned rats had an increase in CW rotations when comparing rats with and without CNO stimulation. This difference approached statistical significance in a one-way ANOVA, F (3, 8) = 6.515, P=0.0153.

After lesioning, rats were tested at 5-weeks with and without CNO. To evaluate our reduction in CCW rotations, we utilized a one-tailed paired samples t-test to evaluate the effect of CNO at 5 weeks between the post lesion without CNO (M = 1743.0 ± 480.5) compared to post lesion with CNO (M = 895.3 ± 801.1), with a mean paired difference of 847.7 ± 511.5. This difference demonstrated a statistical significance reduction in CCW rotations at the five week time point (t(2) = 2.9, p = 0.05). The one-sided 95% confidence interval suggests the true mean difference exceeds –14.7. This study indicates that expression of GAD65 in subthalamic neurons is capable of reducing 6OHDA lesion induced rotations and reduce overall movement disorder in a hemiparkinsonism model.

## Discussion

For our study, we successfully excited and modulated the STN to allow glutamatergic neurons to co-express and release GABA and glutamate(14). Such neurons exist naturally within our nervous system with many vesicles expressing both the vGlut and vGat transporter(15-19). Co-release of GABA by glutamatergic neurons has previously been shown to reduce movement disorders in Parkinson’s rats as well as reduced pain transmission. Co-release of both has been shown to induce biphasic post-synaptic currents and to reduce the activation of AMPA receptors but not NMDA receptors to reduce excitatory currents(20,21). Our study’s findings demonstrated a therapeutic augmentation of parkinsonian phenotype induced in rat models. This was evident by our 5-week time point demonstrating a statistically significant reduction in CCW rotations as well as an increase in CW rotations.

Our study’s findings demonstrated a therapeutic augmentation of parkinsonian phenotype induced in rat models. This was similar to results obtained from Lou et al., 2002, where co-expression of GAD65 within the STN reduced rotational behavior after 6OHDA hemiparkinson lesions. Our study was slightly different from the previous studies in 2 ways. First, we selectively targeted subthalamic neuron collaterals extending into the SNr using Cre-dependent intersectional genetics. Second, we co-expressed excitatory DREADDs with the GAD65 to increase neurotransmitter release in the presence of CNO. In this study, we observed a statistically significant reduction in CCW rotations as well as an increase in CW rotations in 6OHDA lesioned rats treated with GAD65+hM3dq after CNO application, but not in the absence of CNO. We originally expected transduction with GAD65 to reduce rotations alone, and the activation of hM3Dq by CNO would then further reduce them. However, anatomically we observe much fewer subthalamic neurons expressing GAD65 than the Lou paper who directly injected into the subthalamic area. We think that due to the low neuronal labeling, higher levels of stimulation were required to induce GABA release. Additionally, the mechanism by which GABA is released is unknown.

Transduction of glutamatergic sensory neurons demonstrate they reduce pain by releasing GABA through a non-vesicular mechanism, potentially binding to extrasynaptic GABA receptors(14).

Our pre-lesion data was a bit confusing due to the nature of the rotation preference being in opposite direction (CCW) when treated with CNO. Rotational preference should be in the opposite direction if GABA suppressed glutamate induce excitability of striatal neurons. While this may seem counterintuitive, the basal ganglia are a network of excitatory and inhibitory circuits that requires a balance amongst all signals. Additionally, some neurons within the subthalamus expressed only hM3Dq and after activation with CNO might increase excitability slightly under normal conditions. Despite the pre-lesioned rats lacking a parkinsonian phenotype, our therapeutics were previously injected to the lesion and are able to be actively stimulated by the CNO causing an imbalance in the basal ganglia pathway leading to motor deficits. Nevertheless, this data overall indicates an active pathway between the STN and the SNr that we accurately targeted and augmented via DREADDs and GAD65.

The primary limitation of our study was our small sample size. With only three adult rats/group representing our statistically significant data, our statistical power was diminished. This sample size issue was multifactorial but mostly constrained by rat death. Rat death can be attributed to complications with surgery, anesthesia, infection and caging. Other limitations in our study included our PD lesioning model and our behavior assessment model. PD phenotype induction via 6-OHDA injection into the medial forebrain bundle is established as one of the first PD lesioning model. It works by rapidly oxidizing and producing reactive oxygen species to cause mitochondrial dysfunction and subsequent neuron degeneration. Despite 6-OHDA lesioning being a hallmark in the field, it has limitations including its acute effect, lack of blood brain barrier protection, and lack of Lewy Body formation (22). PD pathophysiology in humans contradicts this model as it is gradual in onset with progressive neurodegeneration and Lewy Body formation. Regarding our behavior model, amphetamine induced rotations is a recognized route to evaluate PD behavior (23). Nevertheless, in this model, analysis of rotation quantity and direction characterizes PD phenotype and recovery of function as gross motor function. Unfortunately, while gross motor function is affected in humans with PD, PD has further physical manifestations such has bradykinesia, postural instability, tremor, etc, that are unable to be fully represented by this model. Considering these differences, translation of these results into human models are constrained.

## Conclusions

Findings elicited in this study contributes to PD research and neuroscience as a whole. Our confirmation of the neural network between the STN and SNr lends an important source of manipulation for further advancements in basal ganglia research across all disease states. Therapies like these also have a large implication in the future of PD management; more specifically, these findings can greatly impact patients with medically refractory PD. Minimally invasive injections demonstrated in this research may serve as an essential alternative for high-risk surgical candidates as well as surgically apprehensive patients.

## Author contribution

T.C. constructed the GAD65 plasmid and generated all AAVs used.

N.S., J.R., S.P. performed the experiments or analyzed the behavioral data. N.S., R.S., G.S. designed the experiments and wrote the paper.

## Acknowledgements

This work was partially funded by the Summer Research Program from Lewis Katz School of Medicine at Temple University.

## Materials and Methods

### Plasmid construction and virus production

To induce GABA synthesis, we used an adeno-associated viral vector to express GAD65 and eGFP. The GAD65 transcript was generated by PCR from pAAV-EF1a-double floxed-Gad65-WPRE-HGHpA (13)(Addgene #184631). A polycistronic cassette encoding GAD65 and eGFP linked by a P2A cleavage peptide were subcloned into an AAV DIO backbone containing the human synapsin 1 (hSyn) promoter, the woodchuck posttranscriptional regulatory element (WPRE) and the bovine growth hormone polyA (bGH). This plasmid was sequenced to verify integrity of inserts were all in frame and without mutation. To induce excitation of the STN-SNr pathway, we used an adeno-associated viral vector expressing excitatory DREADDs (hM3Dq) and mCherry. A polyscistronic cassette (pAAV-hSyn-DIO-hM3D(Gq)-mCherry) was given as a gift from Bryan Roth; Addgene plasmid #44361. Recombinant AAV was produced in HEK293T cells using transient triple transfection with rep/cap, adenoviral helper, and AAV transgene plasmids. 72 hours post-transfection, cells were collected and lysed by freeze/thaw cycling and sonication. Viral particles were concentrated using PEG precipitation and purified using two rounds of cesium chloride ultracentrifugation. Virus concentration was determined using SYBR Green qPCR amplifying the AAV ITRs. Viral titers were as follows: AAV2-retro-hSyn-DIO-hM3Dq-mCherry, 1.72×1013 GC/mL; AAV2-scCre, 2.97×1012 GC/mL; AAV2-retro-hSyn-DIO-GAD65-P2A-eGFP, 7.0×1013 GC/mL.

### Surgical procedures

To test our hypothesis, we conducted an in vivo animal study using the 9 Sprague Dawley rats.

This study was approved by the Temple Institutional Animal Care and Use Committee and the University Lab and Animal Resource Committee. Adult rats were appropriately anesthetized and underwent a stereotactic craniectomy of the left skull. Viruses containing GAD and/or DREADDs were injected with a needle unilaterally into the SNr and Cre recombinase was injected into the STN (Fig. 1). With this model, therapies injected into the SNr travelled retrograde to the STN where Cre recombinase was injected via an adeno-associated viral vector (AAV2-scCre); therefore, the protein expression is localized to target STN projections to the SNr specifically. Rats were then left to recover for 10 days before undergoing a second0020surgery to induce the HemiParkinsonian-like phenotype. This was carried out by injecting rats with 6-hydroxydopamine (6-OHDA) into the left medial forebrain bundle ipsilateral to the previous injections to destroy dopamine neurons in the SNc (Fig. 1).

Sprague-Dawley rats were anesthetically induced using inhaled isoflurane with oxygen. Rats were then properly anesthetized using a xylazine:ketamine:saline solution mixed 1:1:2 that was injected intraperitoneally with 1/1000 of their body weight grams to ml (300g rat injected with 0.3 ml). Rat heads were then prepped using betadine and ethanol. All injection coordinates were found using a rat brain atlas (7), with the incisor bar positioned 3.3mm below the horizontal zero, as specified in atlas. For therapeutic injections into the SNr, 0.5 µl of mixed 1:1 saline solution of retro-AAV2-DIO-hM3Dq-mCherry:retro-AAV2-DIO-GAD-GFP was injected unilaterally in two locations into the SNr of rats using the following stereotaxic coordinates: -5.52 AP (from bregma), -1.8 ML, +8.2 DV (from brain surface) and -5.52 AP (from bregma), -2.5 ML, +7.6 DV (from brain surface). Vector was infused at the rate of 0.1 µl/min, and the needle was left in situ for an additional 10 min. For therapeutic injections into the STN, 0.2 µl of saline solution containing AAV2-Cre was injected once unilaterally into the STN of the same rat using the following stereotaxic coordinates from bregma: -3.84 AP, -2.3 ML, and +8.2 DV (from brain surface). Surgery sites were stapled close and rats were given Saline (2 ml) and Rimadyl (0.5 ml) injected subcutaneously for post op pain.

### 6-OHDA Lesioning

6-OHDA was mixed 3 µg/1 µl with 0.1% ascorbic acid. 2.5 µl of the solution was injected at two depths unilaterally into the left medial forebrain bundle using the following coordinates: –2.28 AP (from bregma), -1.9 ML, and +8.1 and +8.8 DV (from brain surface) with the incisor bar placed at 3.3 mm below the horizontal zero. 6-OHDA was injected at the rate of 0.25 µl/min for and the needle was left in situ for an additional 10 min.

### Behavior

Behavior was elicited with and without CNO stimulation via intraperitoneal injection. CNO was prepared in saline solution and rats were dosed 4mg/kg 20 minutes prior to start. For the amphetamine-induce rotation assessment, rats were injected intraperitoneally with amphetamine in saline solution at 5mg/kg. Rats were harnessed and attached to a rotation counter while being placed in a large bowl. Rat rotations quantified as CCW vs CW were counted over the course of 90 minutes. Behavior was analyzed with and without CNO stimulation at two time points: pre-lesional and post-lesional 5 weeks. This assessment entailed rats being injected with amphetamine, harnessed and attached to a rope with a spin counter, then selectively stimulated with CNO to determine the direction and amount of spins recorded. Selective CNO ligand stimulation of DREADDs at each time point allowed each rat to serve as their own control being as the therapies would only be activated after CNO stimulation. Behavioral assessments were statistically evaluated via T-tests. After surgeries and behavioral assessment rats were euthanized and perfused for immunohistochemistry. To ensure appropriate targeting of the basal ganglia and communicating pathways, we used immunofluorescence which depicted appropriate staining of projections from the STN to SNr. Prior to procedure, precursor rats were trialed for appropriate injection location and quantity.

### Immunological and Inflammatory Responses

At 5 weeks post-injection, subgroups of rats were deeply anaesthetized with Euthasol and perfused intracardially with 0.01M phosphate-saline buffer followed by 4% paraformaldehyde. The brain was removed and placed into 4% paraformaldehyde solution overnight and then transferred to 30% sucrose solution for 72 hours. Serial coronal 30µm sections were cut at –20°C using a freezing cryostat (Leica, Germany) through the subthalamic and nigral serial levels. Sections were blocked with 5% donkey serum and 1% Triton X-100 before been incubated with overnight with primary antibody diluted in PBS containing 5% serum. The primary antibodies and the dilutions were Rabbit anti-dsRed (1:500; Takara Bio) and Goat anti-GFP (1:500; Rockland Immunochemicals). Secondary antibodies were Donkey anti-Rabbit AF594 (1:800; Jackson ImmunoResearch), and Donkley anti-Goat AF488 (1:800; Jackson ImmunoResearch).

